# From sea surface to seafloor: a benthic allochthonous eDNA survey for the abyssal ocean

**DOI:** 10.1101/2020.05.07.082602

**Authors:** Olivier Laroche, Oliver Kersten, Craig R. Smith, Erica Goetze

**Affiliations:** Department of Oceanography, School of Ocean and Earth Science and Technology, University of Hawaii at Manoa, Honolulu, USA; Centre for Ecological and Evolutionary Synthesis, Department of Biosciences, University of Oslo, Norway

**Keywords:** environmental DNA, metabarcoding, legacy eDNA, deep sea, abyssal plains, Clarion Clipperton Zone (CCZ), deep-sea mining

## Abstract

Diverse and remote deep-sea communities are critically under-sampled and increasingly threatened by anthropogenic impacts. Environmental DNA (eDNA) metabarcoding could facilitate rapid and comprehensive biotic surveys in the deep ocean, yet many aspects of the sources and distribution of eDNA in the deep sea are still poorly understood. In order to examine the influence of the water column on benthic eDNA surveys in regions targeted for deep-sea polymetallic nodule mining, we investigated the occurrence of pelagic eDNA across: (1) two different deep-sea habitat types, abyssal plains and seamounts, (2) benthic sample types, including nodules, sediment, and seawater within the benthic boundary layer (BBL), and (3) sediment depth horizons (0-2 cm, 3-5 cm). Little difference was observed between seamounts and the adjacent abyssal plains in the proportion of legacy pelagic eDNA sampled in the benthos, despite an > 1000 m depth difference for these habitats. In terms of both reads and amplicon sequence variants (ASVs), pelagic eDNA was minimal within sediment and nodule samples (< 2%), and is unlikely to affect benthic surveys that monitor resident organisms at the deep seafloor. However, pelagic eDNA was substantial within the BBL (up to 13 % ASVs, 86% reads), deriving both from the high biomass upper ocean as well as deep pelagic residents. While most pelagic eDNA found in sediments and on nodules could be sourced from the epipelagic for metazoans, protist legacy eDNA sampled on these substrates appeared to originate across a range of depths in the water column. Some evidence of eDNA degradation across a vertical sediment profile was observed for protists, with higher diversity in the 0-2 cm layer and a significantly lower proportion of legacy pelagic eDNA in deeper sediments (3-5 cm). Study-wide, our estimated metazoan sampling coverage ranged from 40% to 74%, despite relatively large sample size. Future deep-sea eDNA surveys should examine oceanographic influences on eDNA transport and residence times, consider habitat heterogeneity at a range of spatial scales in the abyss, and aim to process large amounts of material per sample (with replication) in order to increase the sampling coverage in these diverse deep ocean communities.

## 1 | Introduction

Deep sea ecosystems are facing increasing pressure from anthropogenic activities, including climate change and future seabed mining (Boschen, Rowden, Clark, & Gardner, 2013; Halpern et al., 2015; Sweetman et al., 2017). The deep sea, defined as the ocean and seafloor below 200 m, represents the largest habitat on our planet, covering ~ 65 % of Earth’s surface (Thistle, 2003; Thurber et al., 2014). The deep ocean is a significant regulator of carbon sequestration and nutrient regeneration, and provides habitat and trophic support to a multitude of organisms (Le, Levin, & Carson, 2017). Characterizing and monitoring the health of deep ocean ecosystems is important, given their role in maintaining Earth’s systems.

The first large-scale deep-sea mining is likely to occur in the Clarion Clipperton Zone (CCZ), a region of particularly valuable mineral resources (nickel- and copper-rich manganese nodules) in the abyssal equatorial Pacific (Wedding et al. 2013, 2015). At > 3 million km^2^, the CCZ spans a wide range of physical and biological gradients, including polymetallic nodule density, particulate organic carbon flux and bathymetry, and is host to diverse biological communities (Glover et al. 2002, Smith et al. 2008a, Wedding et al. 2013). Approximately 30% of the CCZ management area has been allocated to exploration nodule mining through 16contracts granted by the International Seabed Authority (ISA). Benthic communities in the CCZ are typically characterized by high biodiversity and low biomass (Ramirez-Llodra et al., 2010, Kaiser et al, 2017, Smith et al. 2019), due to severe food limitation at abyssal depths (Smith et al. 2008a). A high proportion of species are rare, rendering adequate sampling coverage particularly difficult to achieve (Smith et al., 2008a; Simon-Lledó et al., 2019; De Smet et al., 2017). Regional species diversity is enhanced by habitat heterogeneity at a range of spatial scales, including the presence of polymetallic nodules that serve as hard substrates in a predominantly soft sediment ecosystem (Amon et al., 2016; De Smet et al., 2017; Kaiser, Smith, & Arbizu, 2017; Simon-Lledó et al., 2019; Vanreusel, Hilario, Ribeiro, Menot, & Arbizu, 2016), as well as deep seamounts (relief > 1,000 m above the surrounding seafloor; Clark et al., 2010) that may alter nutrient and particle fluxes and influence associated biological communities (Genin & Dower, 2007; Lavelle & Mohn, 2010; Rowden, Dower, Schlacher, Consalvey, & Clark, 2010; Samadi, Schlacher, & de Forges, 2007). Biodiversity and species ranges remain poorly characterized across the CCZ, making predictions of the impacts of large-scale mining problematic.

Environmental DNA (eDNA) surveys could be particularly valuable for monitoring community change and environmental impacts in the deep sea, given high biodiversity, limited taxonomic descriptions of the fauna, and the challenges of sampling in remote deep ocean habitats (Boschen et al. 2016, Sinniger et al. 2016; Kersten et al., 2019, Laroche et al. in revision). eDNA metabarcoding is a sensitive and cost-efficient tool for biodiversity assessment (Goodwin et al., 2017; Seymour, 2019), with advantages over visual or whole-animal surveys in the ability to capture the hidden diversity of cryptic microbial eukaryotes and to indirectly detect the presence of recently living organisms through cellular debris or extra-cellular DNA (Deiner et al., 2017; Kelly, 2016). However, several methodological aspects regarding eDNA surveys in the deep ocean remain unresolved, including the extent of legacy DNA (extracellular or non-living material) sinking with detrital particles from overlying pelagic ecosystems that arrives in the abyss (*but see* Pawlowski et al., 2011; Morard et al., 2017 for foraminifera). eDNA degradation rates also greatly influence organismal detection; previous studies have reported eDNA half-life ranging from 10 to 50 hours, and complete turnover from a few hours to several months across a range of environmental conditions (see Barnes et al., 2014; Collins et al., 2018). Both abiotic and biotic factors, including temperature, pH, oxygen concentration and microbial activity, have been found to strongly influence eDNA persistence (Armbrecht et al., 2019; Barnes & Turner, 2016; Seymour et al., 2018; Strickler, Fremier, & Goldberg, 2015). eDNA adsorption to sediment particles strongly protects it against hydrolysis and DNAse activity (Torti, Lever, & Jørgensen, 2015), with low temperatures in the deep sea favorable to DNA preservation (Corinaldesi et al. 2008). Because eDNA may be better preserved in deep sea sediments than in seawater, sample substrates from benthic boundary layer (BBL) seawater, polymetallic nodules, and sediments may integrate eDNA inputs over different timescales.

We used an eDNA metabarcoding approach to assess the influence of allochthonous pelagic eDNA on biodiversity surveys at the abyssal seafloor in the western CCZ (Fig. 1). Specifically, our objectives were to evaluate pelagic eDNA occurrence in different: (1) deep sea habitats, including abyssal plains/hills versus seamounts, (2) benthic sample types, including polymetallic nodules, sediment, and seawater within the BBL, and (3) sediment horizons (0-2 cm, 3-5 cm). We also evaluate methods for eDNA recovery from diverse sample substrates and the sampling effort required to fully capture diverse assemblages in the abyssal CCZ. We hypothesize that: (1) most pelagic eDNA recovered from the benthos will have originated in the epipelagic, as production and biomass are highest within this depth zone, (2) the proportion of eDNA shared between pelagic and abyssal samples (BBL seawater, sediment, nodules) will be highest within BBL seawater, rather than in sediments or on nodules, as a range of species may inhabit both the deep pelagic and epi-benthic zones (e.g. meroplankton; Kersten et al., 2017), and (3) pelagic eDNA will occur in higher proportions within the upper sediment horizon (0–2 cm) compared to deeper horizons (3–5 cm) due to detrital eDNA settlement on the surface layer and DNA degradation over time within deeper sediments.

**Fig. 1.**
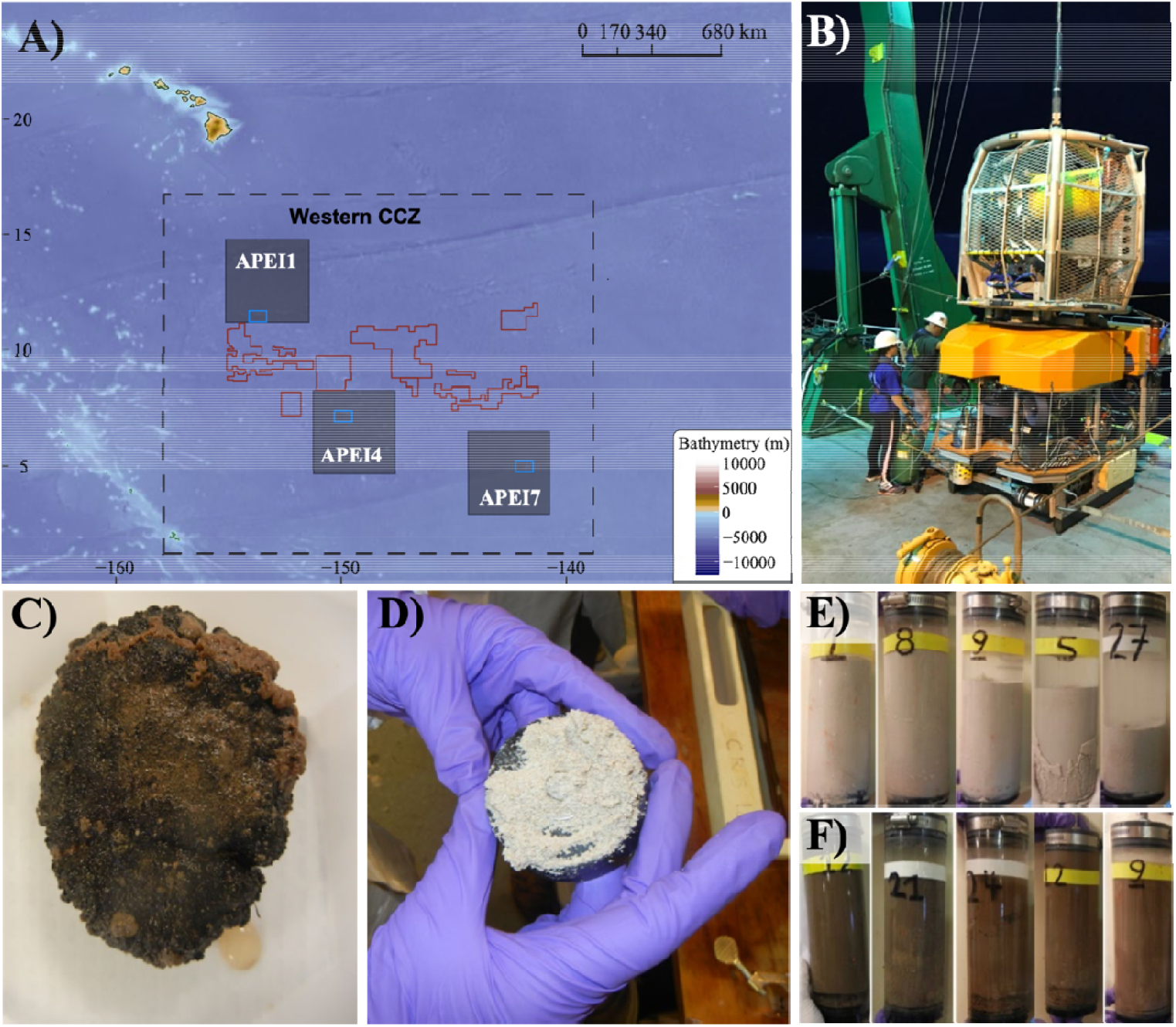
Sampling sites, sampling tools (ROV Lu’ukai), and substrates collected. (A) Map of the study areas within the western Clarion Clipperton Zone (Western CCZ), with the locations of Areas of Particular Environmental Interest (APEIs 1, 4, 7) indicated (grey squares). Sampling regions within each APEI are marked by blue rectangles. Exploration mining contractor (red outline) areas within the western CCZ are also shown. (B) Remotely operated vehicle (ROV) Lu’ukai, used for collection of sediment and polymetallic nodules. (C) Example of a polymetallic nodule. (D) Example of seamount sediment texture, largely made up on foraminiferal tests. A thin slab of sediment is shown on top of a black rubber plate (push core base). (E) Seamount sediment cores. F) Abyssal plain sediment cores.

## 2 | Materials and methods

### 2.1 | Sample collection

Samples were collected in Areas of Particular Environmental Interest (APEIs) 1, 4 and 7 of the western CCZ (Fig. 1; Tables 1, 2, S1), with one seamount and the adjacent abyssal plain/hills sampled within each APEI. APEIs are seafloor areas currently closed by the International Seabed Authority to seabed mining (Wedding et al., 2013). Seamount summit depths ranged from 3100m (APEI 7) to 3900m (APEI 1) and were all > 1000 m above the surrounding seafloor. Adjacent abyssal plain/hill sites were sampled over 15 km away from the seamount ridgeline (APEI 7) or base (APEI 4/APEI 1) to reduce the effect of seamount processes on the abyssal habitats sampled. Due to ROV technical constraints, only seawater samples were collected over the seamount in APEI 1.

**Table 1.** Station overview for ROV dives from which sediment and nodules were collected for eDNA.

| Station # | APEI | Habitat | ROV dive # | Date | Latitude | Longitude | Water depth (m) |
| --- | --- | --- | --- | --- | --- | --- | --- |
| KM1808-25 | 7 | Plain | 4 | 5/25/18 | 5° 6.9271 N | 141° 53.8015 W | 4860 |
| KM1808-33 | 7 | Seamount | 6 | 5/27/18 | 4° 53.28 N | 141° 45.4299 W | 3133 |
| KM1808-36 | 7 | Plain | 7 | 5/28/18 | 5° 3.6190 N | 141° 49.8836 W | 4874 |
| KM1808-43 | 4 | Plain | 8* | 6/01/18 | 7° 2.1617 N | 149° 56.3686 W | 5040 |
| KM1808-47 | 4 | Plain | 9* | 6/2/18 | 6° 59.2646 N | 149° 54.7421 W | 5019 |
| KM1808-49 | 4 | Seamount | 10 | 6/5/18 | 7° 16.2918 N | 149° 46.7055 W | 3562 |
| KM1808-61 | 4 | Plain | 12* | 6/6/18 | 6° 59.2865 N | 149° 55.9757 W | 5009 |
| KM1808-66 | 1 | Plain | 13* | 6/10/18 | 11° 16.5 N | 153° 44.64 W | 5240 |
| KM1808-70 | 1 | Plain | 14* | 6/11/18 | 11° 15.12 N | 153° 36.37 W | 5206 |

ROV dives from which nodules could be collected are indicated by a star.

Sediment samples were obtained with the ROV Lu’ukai using 7-cm diameter push cores. Typical ROV dives yielded two cores in close proximity (meters) near the start of the dive and another 2 cores at mid-dive, on average 2.3 km away from the starting point. Cores were subsampled for eDNA at 0-2 cm and 3-5 cm sediment horizons using single-use sterile syringes (60 mL) to extract one mini-core from each sediment horizon. Samples were cryopreserved at −80 °C until processing. Between ROV dives (Table 1), sediment-processing gear and push-core tubes were treated with 10% bleach and rinsed with ddH_2_O to prevent DNA contamination. Subsampling equipment (core slicer, guide) was rinsed in ddH_2_O between cores from the same ROV dive. For APEI 1, two replicate subsamples were taken from each sediment horizon and core. Polymetallic nodules were collected from the same ROV dives using either push cores or the manipulator arm of the ROV, with nodules placed into the BioBox sample holder on the ROV, which was sealed during ROV recovery from the seafloor. Upon arrival on shipboard, nodules were transferred to sterile whirl-pack bags and cryopreserved (−80 °C).

Seawater samples were collected using conductivity-temperature-depth (CTD) casts (Table 2) with a 24 Niskin bottle rosette (10L). Four CTD casts per APEI were conducted, two over the abyssal plain/hills and two over the seamount (Table 2). Seawater was collected at seven depths within the water column: 5 meters above bottom (mab), 50 mab, bathypelagic depths (3000 m over plains/hills, 2500 or 2000 m over seamounts, depending on summit depth), in the deep mesopelagic at 1000 m, mesopelagic at 500 m, deep chlorophyll maximum (DCM; between 90 m and 60 m), and at 5 m depth in near surface waters. Eukaryotic productivity and biomass vary greatly across depth in the water column, with exponential declines in pelagic animal biomass (e.g., Vinogradov 1970, Angel & Baker 1982). As a result, while it is common for eDNA studies to sample the epipelagic community with small seawater volumes (e.g., 250 mL to 2 L liters, Günther et al. 2018; Jeunen et al., 2019; Shulse et al., 2017; Sigsgaard et al., 2017; Thomsen et al. 2012), larger volumes are required to offset expected declines in eDNA concentration and adequately sample communities at depth (e.g., as in Shulse et al. 2017). Therefore, filtered seawater volumes varied across depth in this study, with 5 L per replicate sampled at 5 mab, 50 mab and bathypelagic depths, 4 L filtered in the deep mesopelagic (1000m), 2 L in the mesopelagic (500m), and 1 L filtered per replicate at the DCM and in the near surface. Four to six replicates were taken from each CTD cast and depth. To assess and eliminate cross-contamination, negative controls (double-distilled water; ddH_2_O) were collected for each CTD cast (1 L for each of 2 replicates). eDNA was collected on sterile 0.2 μm Supor filters (Pall) using 47 mm inline polycarbonate filter holders and two peristaltic pumps. Filters were preserved in 1 ml of RNALater (Invitrogen), flash frozen in liquid nitrogen, and held at −80°C until processing. Between CTD casts, the workspace and all sampling equipment were treated with 10% bleach for a minimum of 30 minutes, followed by three ddH_2_O and three seawater rinses. To avoid contamination during sample collection, personal protective equipment included disposable lab coats and nitrile gloves for all involved personnel. Table S1 reports all sediment, nodule and seawater samples included in this study.

**Table 2.** List of conductivity, temperature and depth (CTD) casts from which seawater was collected for eDNA. Max depth (m) indicates the seafloor depth at the sampling site.

| Station # | APEI | Habitat | Cast # | Date | Latitude | Longitude | Max. depth (m) |
| --- | --- | --- | --- | --- | --- | --- | --- |
| KM1808-12 | 7 | Plain | 1 | 5/22/18 | 5° 3.6252 N | 141° 49.8395 W | 4861 |
| KM1808-13 | 7 | Plain | 2 | 5/22/18 | 5° 3.2156 N | 141° 55.5066 W | 4850 |
| KM1808-27 | 7 | Seamount | 4 | 5/25/18 | 4° 53.6165 N | 141° 43.7357 W | 3228 |
| KM1808-35 | 7 | Seamount | 6 | 5/28/18 | 4° 53.310 N | 141° 46.1472 W | 3060 |
| KM1808-48 | 4 | Plain | 7 | 6/3/18 | 6° 58.8754 N | 149° 55.6328 W | 4994 |
| KM1808-54 | 4 | Seamount | 8 | 6/4/18 | 7° 17.3894 N | 149° 51.2256 W | 3919 |
| KM1808-56 | 4 | Seamount | 9 | 6/5/18 | 7° 15.8752 N | 149° 46.4703 W | 3560 |
| KM1808-60 | 4 | Plain | 10 | 6/6/18 | 7° 02.2456 N | 149° 56.3370 W | 5034 |
| KM1808-67 | 1 | Plain | 11 | 6/10/18 | 11° 16.4768 N | 153° 44.7812 W | 5232 |
| KM1808-71 | 1 | Plain | 12 | 6/11/18 | 11° 15.1408 N | 153° 36.3776 W | 5196 |
| KM1808-76 | 1 | Seamount | 13 | 6/12/18 | 11° 29.6389 N | 153° 36.9525 W | 4045 |
| KM1808-77 | 1 | Seamount | 14 | 6/12/18 | 11° 29.5117 N | 153° 39.0729 W | 4183 |

Filtered seawater volume per replicate sample and per depth was 5 L at 5 meter above bottom (mab), 50 mab and at bathypelagic (2000-2500 m for seamounts and 3000 m for plains), 4 L at deep mesopelagic (1000 m), 2 L at mesopelagic (500 m), and 1 L at the DCM and near surface (5 m) samples.

### 2.2 | DNA extraction

Sediment samples were homogeneized with a sterile metallic spatula, subsampled for 10 g, and processed with the PowerMax^®^ Soil kit (QIAGEN, California, USA) following the manufacturer’s protocol. Purified eDNA was eluted in 1 mL of ddH_2_O. Prior to eDNA extraction, polymetallic nodules were ground and homogenized inside their whirl-pack bag using a 16 g ceramic pestle. No detectable quantities of DNA could be obtained using the PowerMax^®^ Soil kit with ~ 10 g of nodule material, likely as a result of very low DNA concentrations on nodules. We therefore used the FastDNA™ Spin kit (following the manufacturer’s protocols), processing ten subsamples of ca. 500 mg for each nodule. Due to low eDNA concentrations per subsample (mean concentration of 0.382 ng/μL), replicates were pooled in pairs and concentrated to ~ 1ng/μL with the DNA Clean & Concentrator kit (Zymo Research, California, USA) to obtain sufficient DNA for polymerase chain reaction (PCR) amplification.

For seawater samples, four extraction protocols were tested in pilot experiments: (1) a Cetyl Trimethyl Ammonium Bromide (CTAB) and ethanol precipitation protocol based on Renshaw et al. (2015), (2) the DNeasy^®^ Blood and Tissue kit (QIAGEN, California, USA) using a modified protocol from Djurhuus et al. (2017), (3) the E.Z.N.A^®^ Water DNA kit (OMEGA, Georgia, USA), and (4) the DNeasy^®^ Plant Mini kit (QIAGEN, California, USA), using amodified protocol based on Pearl et al. (2008) and Shulse et al. (2017). Using biological triplicates, the latter protocol provided the highest DNA yield and best purity measurements, and was therefore used to process all seawater samples (*see* the supplementary material for the full protocol). Due to low DNA concentration in the 5 and 50 mab samples (mean concentration of 0.573 ng/μL), replicates (2 x 5 L each) were pooled together and concentrated to ~ 1ng/μL with the DNA Clean & Concentrator kit (Zymo Research, California, USA) to obtain sufficient DNA for amplification.

To assess and eliminate cross-contamination, an extraction blank was included for each sample substrate type (sediment, nodule, seawater). All sample handling and extraction steps took place in a dedicated laboratory that had never been used for PCR amplification.

### 2.3 | PCR amplification and library preparation

The V4 region of the 18S rRNA gene (approximately 450 base pairs [bp]) was PCR amplified using the eukaryotic forward primer Uni18SF: 5’-AGG GCA AKY CTG GTG CCA GC-3’ and reverse primer Uni18SR: 5’-GRC GGT ATC TRA TCG YCT T-3’ (Zhan et al., 2013). Primers were modified to include Illumina™ overhang adaptors. The choice of optimal 18S rRNA hypervariable regions for biodiversity assessments remains under debate (e.g., *see* Hadziavdic et al., 2014, Tanabe et al., 2016). Here, preliminary tests comparing the V1-V2 region using primers from Fonseca et al. (2010) and the V4 region with primers from (Zhan et al., 2013) found highest taxonomic classification and diversity at the genus level (especially for metazoans) with the latter primer set. This marker and primer set were therefore selected for use in the full study.

PCR reactions consisted of 12.5 μL of MyFi™ Mix (Bioline; hot-start, proof-reading Taq polymerase), 0.5 μL of each primer (10 μM), 0.6 μL of Bovine Serum Albumin (300 μM; BSA; Sigma), 1-3 μL of template DNA (min 2.5 ng/μL per reaction) with ddH_2_O added to reach a total volume of 25 μL. The reaction cycling conditions were: 95°C for 3 min, followed by 35 cycles of 95°C for 15 s, 55°C for 30 s, 72°C for 15 s, with a final extension step at 72°C for 1 min. A negative control (no template DNA) was included with each PCR to ensure an absence of contamination.

Purification and quantification of the amplicons was performed with AMPure^®^ XP beads (Beckman Coulter^®^, Indiana, USA) and a Qubit^®^ Fluorometer (Life Technologies), following the manufacturer’s instructions. Purified amplicons were normalised to 2 ng/μL with ddH_2_O, and submitted to the Advanced Studies in Genomics Proteomics and Bioinformatics (ASGPB) of the University of Hawaii at Manoa (HI, USA) for indexing with the Nextera™ DNA library Prep Kit (Illumina, California, USA). Samples were pooled into two libraries and sequenced on two MiSeq Illumina™ runs using V3 chemistry and paired-end sequencing (2×300 bp). Two blank samples containing ddH_2_O were included during the indexing and sequencing process to assess potential contamination. Sequences are available from the NCBI Sequence Read Archive (SRA) under accession numbers SRR9199590 to SRR9199869.

### 2.4 | Bioinformatic analysis

Demultiplexed samples were denoised with the DADA2 pipeline (Callahan et al., 2016), as implemented in Qiime2 (Bolyen et al., 2019), using the qiime dada2 denoise-paired function and default parameters. The DADA2 pipeline is a sequence quality control process that filters low quality reads based on the maximum expected error value (default=2), detects and removes Illumina amplicon artefacts, filters PhiX reads, and merges paired forward and reverse reads using a default minimum overlapping region of 20 bp. DADA2 is particularly efficient at eliminating spurious sequences from Illumina platforms (Callahan et al., 2016). Prior to merging, forward and reverse reads were truncated at 260 bp and 235 bp, respectively, to remove low quality regions and reduce the number of sequences lost during quality filtering. The DADA2 pipeline detects and removes chimeric sequences using a *de novo* approach. Here, chimera removal also was performed using the consensus approach, in which sequences found to be chimeric in a majority of samples are discarded. Taxonomic assignment was performed by training Qiime2’s naive Bayes classifier (Pedregosa et al., 2012) on a SILVA 18S rRNA database (release 132 clustered at 99 % similarity; Quast et al., 2013) trimmed with Qiime2’s feature-classifier extract-reads function and the V4 primers. For downstream analyses, amplicon sequence variants (ASVs) were used instead of operational taxonomic unit (OTUs) in order to retain the highest possible taxonomic resolution in the data. The complete bioinformatics script is provided in supplementary material.

### 2.5 | Data analysis and statistics

Sequencing depth and recovered diversity per sample were inspected using rarefaction curves with the vegan R package (Oksanen et al., 2018). Rarefaction curves indicated that all but one sample (N-26) had sufficient sequencing depth to capture total amplicon richness within the sample. This sample was excluded from all downstream analyses (Fig. S1). Sequences found in any of the negative controls, including field (ddH_2_O), DNA extraction, and PCR blanks were investigated and removed from the dataset. Sequences unclassified at kingdom level and those originating from non-marine taxa also were discarded (taxa assigned to birds, insects, terrestrial mammal families that likely originate from marine vessels waste). The data was then split into two groups: (1) metazoans, and (2) non-metazoans, primarily comprised of protists from the SAR supergroup (Stramenopiles, Alveolata, and Rhizaria; Adl et al., 2019). Amplicon sequence variant (ASV) richness for each sample type (sediment, nodule and seawater) was estimated using the Chao2 index (per sample), an estimator of asymptotic species richness based on the frequency of rare taxa, as well as with a bootstrap method with 200 replicates as outlined in Chao and Jost (2012). When the Chao2 index is estimated per sample, as in this study, rare taxa correspond to those found in very few samples (rather than those that are rare within each sample). Calculations and visualizations were conducted using the iNEXT (version 2.0.19; Hsieh, Ma, & Chao, 2019) and ggplot2 R packages (Wickham, 2016). The sampling coverage, or sampling completeness, estimator was based on the incidence of singletons and doubletons across samples (defined in equation 4a of Chao & Jost 2012). We used base coverage as a metric for comparison among multiple samples; this metric combines rarefaction and extrapolation, and represents the highest coverage value between minimum extrapolated values and maximum interpolated values (*see* Chao et al., 2014).

Alpha and gamma diversity, reported as ASV richness estimated by the Chao2 index, was also compared across the water column. ASV overlap between sediment layers (0-2, 3-5 cm) and water columns depths within the BBL (5, 50 mab) were investigated with Venn diagrams using the Venny 2.1 program (Oliveros, 2015), in order to evaluate the importance of sampling these distinct depth horizons.

The presence of pelagic eDNA at the seafloor, here defined as ASVs from samples of the near surface to the bathypelagic ocean (0 to 3000 m over abyssal plains, 0 to 2000-2500 m over seamounts, inclusively) that were simultaneously found in benthic samples, was examined from 2 perspectives: 1) the pelagic perspective, representing the proportion of pelagic ASVs and reads concurrently found in the benthic environment, and 2) the benthic perspective, representing the proportion of benthic ASVs and reads for each sample type (BBL seawater, sediments, nodules) that were concurrently found within pelagic samples. The taxonomic composition and proportions of pelagic eDNA found within abyssal habitats were visualized using ggplot2. From the benthic perspective, the proportion of benthic ASVs and reads per sample was also contrasted per sample type and tested for significant difference between habitats and sediment layers with a two-sample Wilcoxon test in R (wilcox.test function), and visualised with box-plots and the ggplot2 R package. To ensure that results were not biased by differences in sampling completeness between habitats, sampling coverage was subsequently conducted (Fig. S2).

## 3 | Results

### 3.1 | High-throughput sequencing

A total of 10,315,003 18S reads were generated (Table S2). Quality filtering, denoising, merging and chimera removal reduced read counts by 54%, leaving an average of 43,523 good quality reads per sample. Removal of sequences found in the blank samples resulted in a mean loss of 2.7 % ASVs per seawater sample and 0.4% of ASVs for nodules (Table S3). Removal of non-marine taxa led to a further loss of 3.3 % ASVs for nodules, 1.5 % for sediment and less than 1 % for seawater samples (Table S3). All subsequent analyses were performed on both filtered and unfiltered data to examine the effect of removing sequences found in blank samples or belonging to non-marine metazoan taxa. However, since this filtering step had no consequences on the results, only analyses performed on trimmed data are presented.

### 3.2 | Sampling coverage and ASV richness

Overall, the achieved sampling coverage was highest for seawater (74 % and 84 % for metazoans and non-metazoans, respectively), followed by nodules (65 % for both groups) and sediment samples (40 % and 64 % for metazoans and non-metazoans, Fig. 2). Gamma diversity, or ASV richness for all samples combined, was highest within sediment samples (1,271 metazoan and 6,107 protist ASVs), followed by seawater (409 and 5,953 ASVs) and nodules (407 and 2,186 ASVs, Fig. 2). Similarly, ASV richness at base coverage was higher in sediment samples than in nodules or seawater samples for both metazoans and non-metazoans (Fig. 2).

**Fig. 2.**
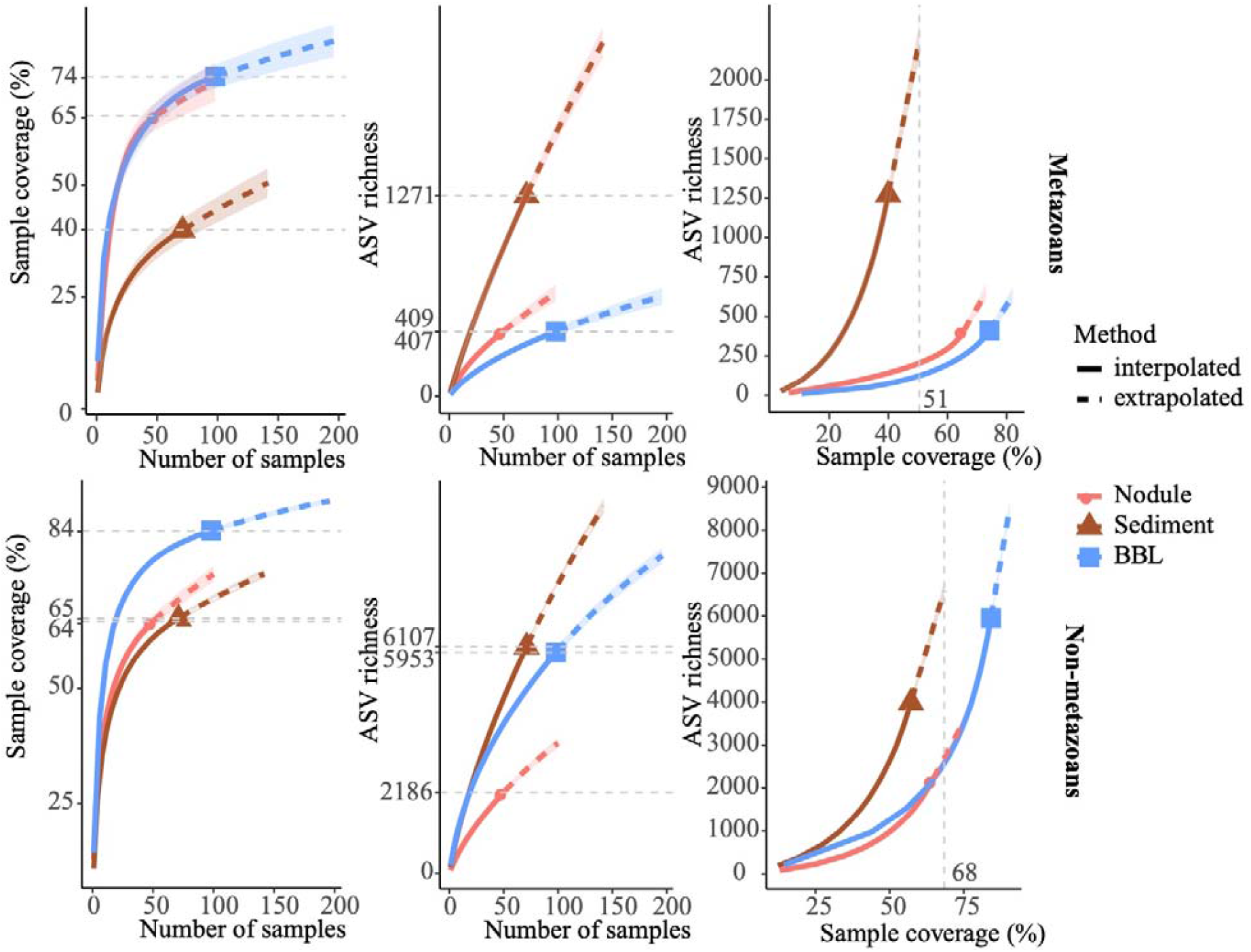
Amplicon sequence variant (ASV) sampling coverage and richness for each sample type (nodule, sediment and seawater). ASV richness was estimated using Chao2 (Chao & Jost 2012). Shaded, coloured areas indicate the 95 % confidence intervals obtained from a bootstrap method based on 200 replicates. Horizontal dotted grey lines (left, center panels) indicate maximum interpolation values for each sample type. Vertical dotted grey lines (right panels) indicate the value at base coverage, defined as the highest coverage value between minimum extrapolated values and maximum interpolated values.

Sampling coverage analyzed at the level of biological replicates indicated that for sediment samples, two replicates of 10 g each led to a mean sampling coverage of 15 % and 39 % for metazoans and non-metazoans of the 0-2 cm layer, respectively, and 8 % and 36 % for the 3-5 cm layer (Fig. S3). Further analysis of community composition in the 0-2 and 3-5 cm layers showed modest ASV overlap between the two (10 % and 19 % of shared ASVs for metazoans and non-metazoans, respectively), with a large fraction of ASVs found solely in the top layer (56 % and 47 % for metazoans and non-metazoans, respectively) (Fig. S4). For nodules, five biological replicates (1 g of material each) extracted from each nodule provided a mean sampling coverage of 58 % for metazoans and 43 % for non-metazoans (Fig. S3). With three biological replicates, sampling coverage in seawater samples averaged 53 % for metazoans and 69 % for non-metazoans across water column depths (Fig. S3). Further analysis of community overlap between the BBL water column depths showed that 5 mab and 50 mab samples shared 23 % and 30 % of metazoan and protist ASVs, respectively, with the 5 mab layer possessing the greater portion of unshared ASVs (48 % and 45 % for metazoans and non-metazoans) (Fig. S4). Overall, 66 % and 73% of total benthic diversity was captured in the upper sediment horizon and the 5 mab BBL seawater sample, respectively.

Mean metazoan ASV richness per sample remained largely uniform across depths in the water column, with slightly higher values in the DCM (12 ± 4) and BBL (5 mab and 50 mab; 14 ± 7) (Fig. 3A). However, sampling coverage varied greatly across water column depths (from 49 % to 73 %), with lowest values or the 5 mab (49%) and 50 mab (57%) samples (Fig. 3B). Significant differences in ASV richness (absence of overlap in confidence intervals) can be observed between 5 mab and the mesopelagic, deep mesopelagic and near surface depths (Fig. 3C). At base coverage (64 %), diversity was highest within the BBL samples (~ 225 and ~ 100 ASVs at 5 mab and 50 mab, respectively), and lowest within the mesopelagic (500m) and deep mesopelagic (1,000 m) samples (~ 55 ASVs; Fig. 3C). A comparison of the sampling effort required at each depth to reach equivalent sampling coverage is shown in Table 3 (at base coverage, 49 %). The water column depths requiring highest sampling effort, or the largest volume of seawater filtered, are 5 mab, 50 mab, the bathypelagic and the DCM.

**Fig. 3.**
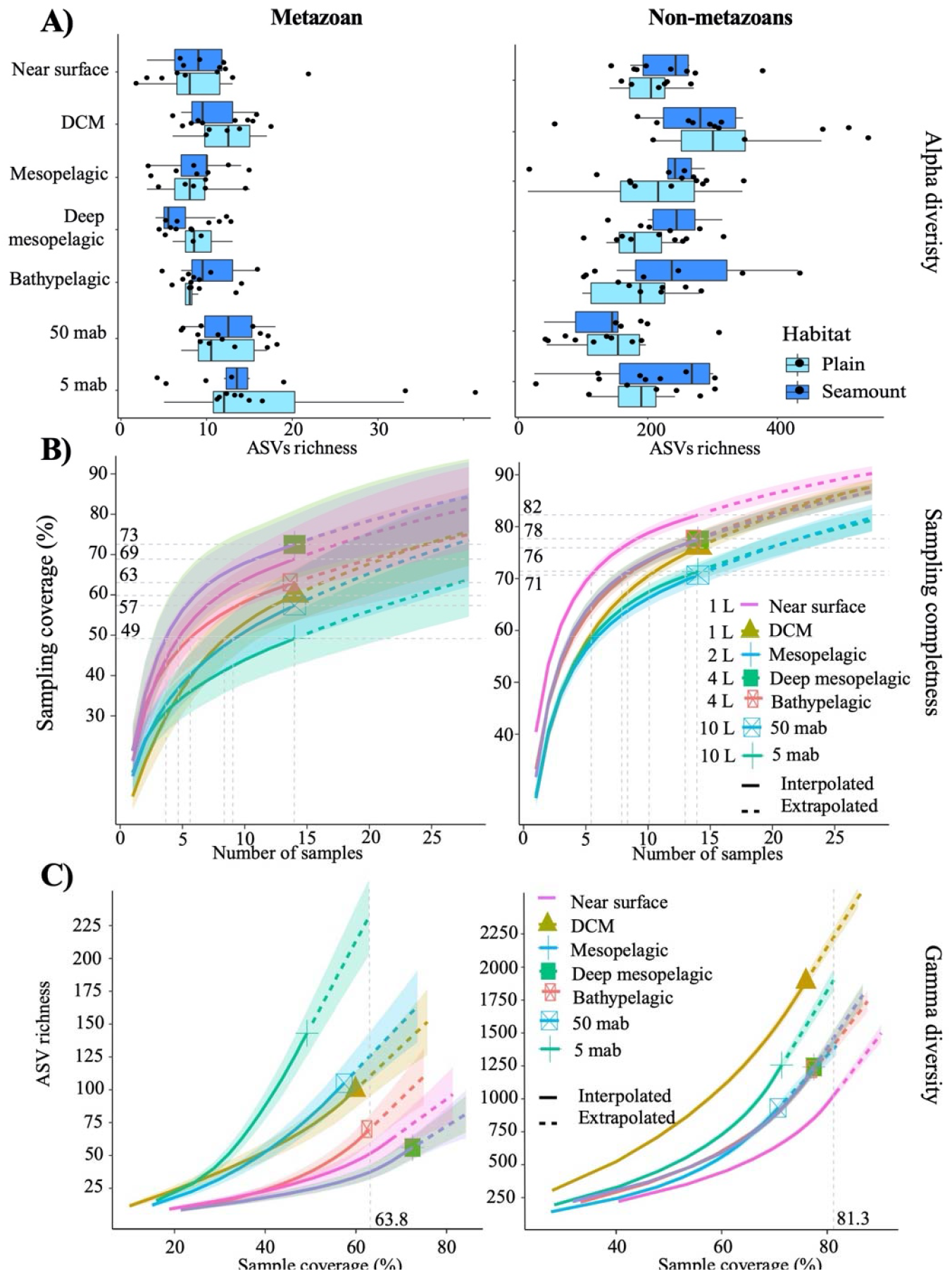
Alpha-diversity, sampling completeness and gamma-diversity across the water column for both metazoans and non-metazoans. (A) Box-plot of mean amplicon sequence variant (ASV) richness observed per sample, within each habitat type (abyssal plains, seamounts). (B) Sampling coverage achieved across the water column, with seawater volumes filtered at each depth as listed. Horizontal grey dotted lines indicate maximum interpolated sampling coverage values for each water column depth horizon. (C) Estimated ASV richness as a function of sampling coverage (Chao2 estimator), across the water column. The vertical dotted grey line represents sample completeness at base coverage, defined as the highest coverage value between minimum extrapolated values and maximum interpolated values. DCM = deep chlorophyll maximum, mab = meters above bottom. For B and C, shaded coloured areas indicate the 95 % confidence intervals obtained by a bootstrap method based on 200 replicates.

**Table 3.**
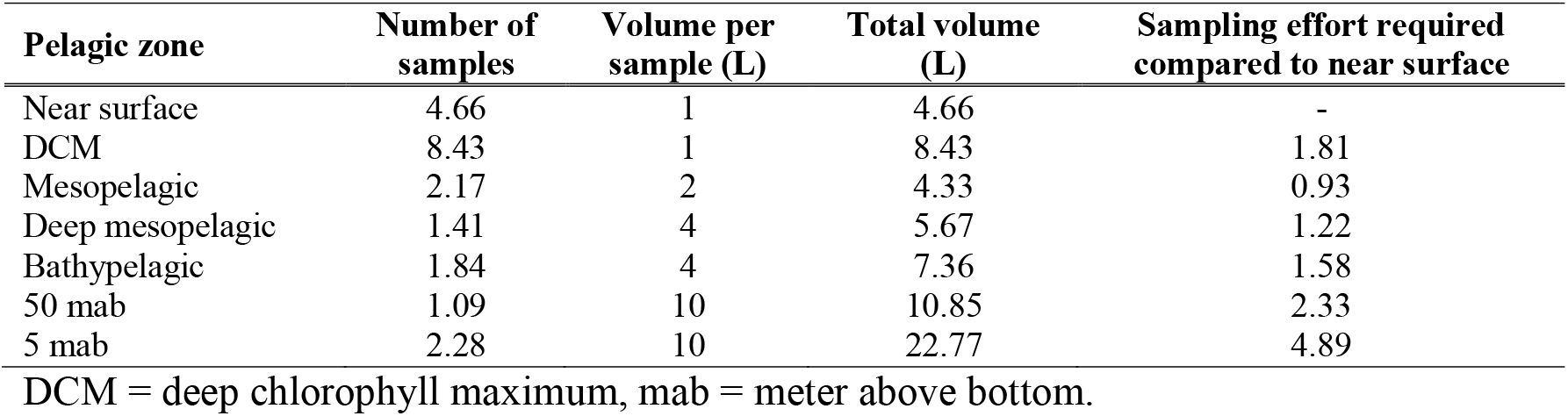
Sampling effort required at each pelagic depth to reach the minimum sampling coverage achieved across depth (49 %) for metazoans.

| **Pelagic zone** | **Number of samples** | **Volume per sample (L)** | **Total volume (L)** | **Sampling effort required compared to near surface** |
| --- | --- | --- | --- | --- |
| Near surface | 4.66 | 1 | 4.66 | - |
| DCM | 8.43 | 1 | 8.43 | 1.81 |
| Mesopelagic | 2.17 | 2 | 4.33 | 0.93 |
| Deep mesopelagic | 1.41 | 4 | 5.67 | 1.22 |
| Bathypelagic | 1.84 | 4 | 7.36 | 1.58 |
| 50 mab | 1.09 | 10 | 10.85 | 2.33 |
| 5 mab | 2.28 | 10 | 22.77 | 4.89 |

DCM = deep chlorophyll maximum, mab = meter above bottom.

For non-metazoans, ASV richness per sample was highest within the DCM (304 ± 131) before gradually declining through the meso- and bathypelagic and then increasing again at 5 mab (Fig. 3A). In contrast to metazoans, non-metazoan sampling coverage remained relatively high and uniform across depths (from 71 % to 82 %; Fig. 3B), although significant differences can be observed between the near surface, BBL and the zones in between at the highest number of samples collected (14). At base coverage, diversity was highest in the DCM (~ 2,200 ASVs) and 5 mab (~ 1,900 ASVs), and lowest at the near surface (~ 1,050 ASVs; Fig. 3C). Comparison of the sampling effort required per depth to reach equivalent sampling coverage (71 %), is shown in Table 4, with 50 mab, 5 mab, deep mesopelagic and DCM requiring the largest volumes of seawater filtered.

**Table 4.** Sampling effort required at each pelagic depth to reach the minimum sampling coverage achieved across depth (71 %) for non-metazoans.

| **Pelagic zone** | **Number of samples** | **Volume per sample (L)** | **Total volume (L)** | **Sampling effort required compared to near surface** |
| --- | --- | --- | --- | --- |
| Near surface | 5.44 | 1 | 5.44 | - |
| DCM | 10.12 | 1 | 10.12 | 1.86 |
| Mesopelagic | 4.44 | 2 | 8.88 | 1.63 |
| Deep mesopelagic | 2.69 | 4 | 10.77 | 1.98 |
| Bathypelagic | 2.33 | 4 | 9.31 | 1.71 |
| 50 mab | 2.29 | 10 | 22.9 | 4.21 |
| 5 mab | 1.51 | 10 | 15.11 | 2.78 |

DCM = deep chlorophyll maximum, mab = meter above bottom.

### 3.3 | Pelagic eDNA in the water column

Metazoan eDNA within the epipelagic derived predominantly from arthropods (mean per sample of 70 ± 13% of reads), followed by chordates (15 ±4 %; essentially composed of appenticularians and thaliaceans) and cnidarians (11 ±4 %) (Fig. 4A). Mesopelagic eDNA differed substantially from the epipelagic in that fewer arthropod reads (< 17 %) and a higher proportion of cnidarian reads (72 %) were observed. The deep mesopelagic and bathypelagic zones contained a high proportion of cnidarian reads (> 75 %), which were mostly unassigned at the family level within the classes Hydrozoa (75 %) and Scyphozoa (4 %), but included reads assigned to families Agalmatidae (9 %), Rhopalonematidae (4 %) and Prayidae (2 %).

**Fig. 4.**
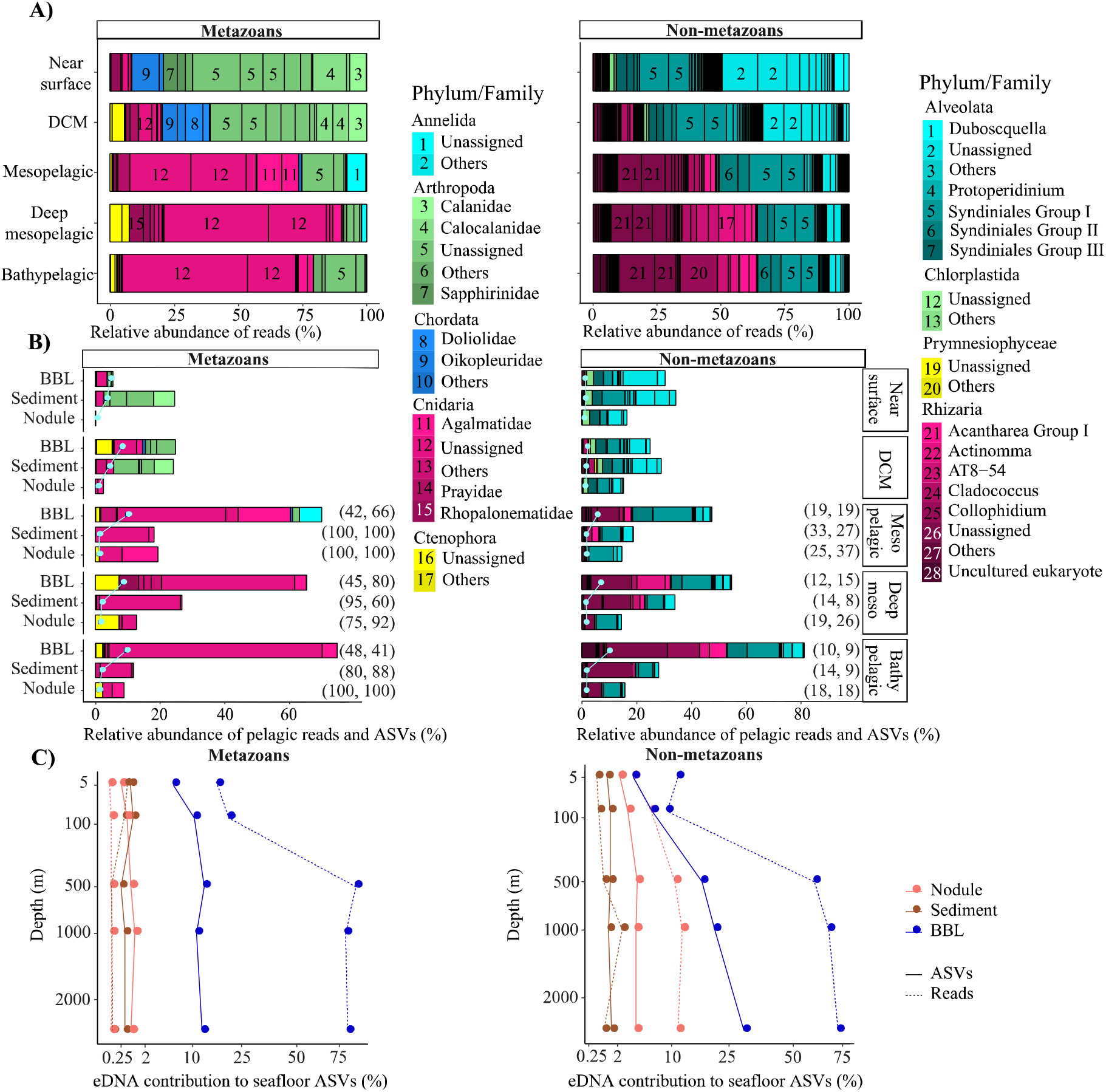
Pelagic eDNA community composition and contribution to diversity at the seafloor. A) Relative read abundance (bars) of phyla and families per pelagic depth zone within each deep ocean habitat (abyssal plains, seamounts). B) Relative abundance of pelagic reads (bars) and pelagic ASVs (pale gray dots and lines) that also were sampled in abyssal sediment, nodule or BBL seawater samples, plotted relative to the diversity sampled in the water column per pelagic depth zone and ocean habitat (pelagic perspective). The percent of pelagic ASVs and associated reads simultaneously found in the epipelagic are indicated on the right side of the bars. C) Proportion of sediment, nodule or BBL seawater ASVs and reads that were found within each pelagic zone sampled. Plot shows the benthic perspective, or pelagic diversity sampled at the seafloor relative to all diversity sampled at the seafloor. BBL = Benthic boundary layer, DCM = deep chlorophyll maximum.

Non-metazoan taxonomic composition also varied substantially across the water column, with Alveolata dominating the epipelagic (84 ± 7 %) and Rhizaria dominating the meso- and bathypelagic (58 ± 10 %; Fig. 4A). Alveolate reads were primarily from Protalveolata, specifically from one of the five Syndiniales groups (31 % of all pelagic reads), as well as the dinoflagellates (22 %). Within the Rhizarians, most sequences belonged to Order Spumellaria (16 % of all pelagic reads), Order Collodaria (9 %), Class Acantharia (4 %) and the RAD B class of the Retaria subphylum (3 %).

### 3.4 | Pelagic eDNA at the seafloor

#### 3.4.1 | Pelagic perspective

A relatively small fraction of the metazoan diversity observed in the upper ocean (near surface, DCM) was sampled at the abyssal seafloor, ranging from 1.2 to 24 % of reads from ASVs that were simultaneously observed in abyssal sediments, nodules or seawater (Fig. 4B). In contrast, a large fraction of the metazoan diversity in the meso- and bathypelagic ocean was found within BBL seawater samples (up to 75 % of reads, 10 % of ASVs; Fig. 4B). Interestingly, the great majority of metazoan pelagic ASVs (> 75 %) and reads (> 60 %) below the DCM that were simultaneously found in sediment and nodules could have been sourced from the epipelagic, as these sequences were also present in the near surface and/or DCM samples.

Many of the patterns observed for metazoans were shared for non-metazoans, including the lower proportion of epipelagic water column diversity sampled at abyssal depths in comparison to meso- and bathypelagic non-metazoan diversity (Fig. 4B). In the mesopelagic and below, the proportion of pelagic protist reads found within BBL seawater samples increased gradually and substantially, from 47 % to 81 % and 6 % to 10 % for reads and ASVs, respectively. In contrast to metazoans, the non-metazoan pelagic ASVs and reads sampled below the DCM that could have been sourced in the epipelagic (also found within near surface and/or DCM samples) was much lower (< 33 % and <37 %, respectively; Fig. 4B). Overall, pelagic protist reads were found in the smallest proportion within nodules (15 ± 1 %), followed by sediments (29 ± 6 %) and seawater samples (48 ± 22 %).

For both metazoans and non-metazoans, the identity of pelagic eDNA found within the benthos closely reflected the overall taxonomic composition of each pelagic zone (Fig. 4A and B), with no evidence of taxonomic filtering in terms of which pelagic taxa were sampled at the abyssal seafloor.

#### 3.4.2 | Benthic perspective

Because nodules and especially sediment samples had higher metazoan ASV richness than seawater (Fig. 2), the proportion of diversity that is pelagic in origin represents a small fraction of the total sampled diversity at the seafloor. For example, pelagic metazoan reads and ASVs corresponded, respectively, to 0.012 % and 0.64 % of the diversity sampled on nodules and 0.15 % and 0.53 % for sediments. However, pelagic eDNA made up a much larger fraction of the BBL eDNA diversity (56 % reads, 13 % ASVs) (Fig. 4C). Within the BBL, most of the pelagic diversity derives from meso- and bathypelagic depths. For metazoans, the percentage of ASVs and associated reads per benthic samples that were simultaneously found in the pelagic were not significantly different between seamount and abyssal plain habitats and upper and lower sediment horizons (Fig. 5), when evaluated as combined pelagic samples of the relevant APEI and habitat (Table S4).

**Fig. 5.**
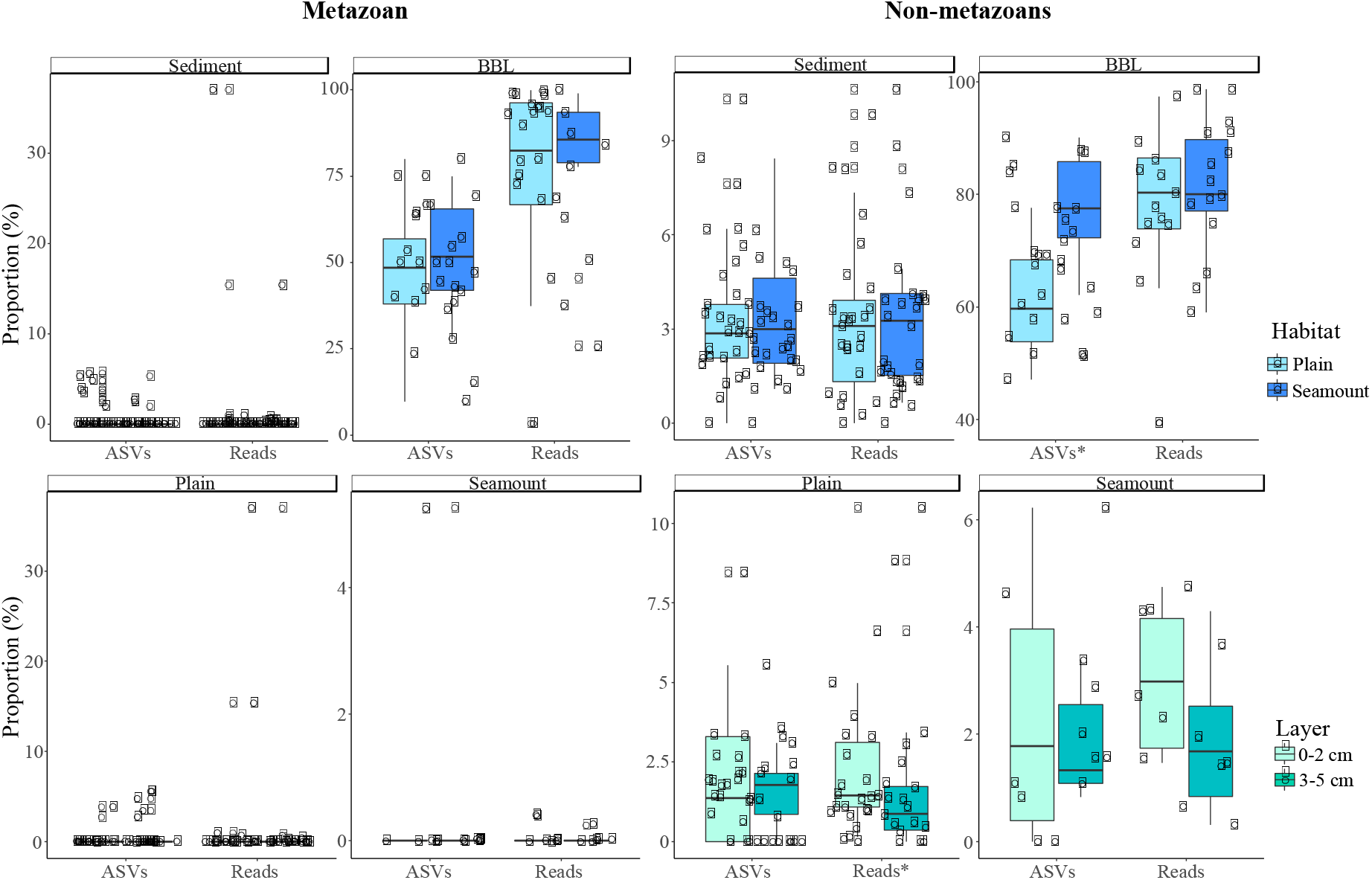
Box-plots of the mean proportion of amplicon sequence variants (ASVs) and their respective reads that derive from pelagic ecosystems (surface to bathypelagic depths) that were sampled at the seafloor across habitats (abyssal plain, seamount) and sediment layers (0-2, 3-5 cm) for both metazoans and non-metazoans. Plot shows the benthic perspective, or pelagic diversity sampled at the seafloor relative to all diversity sampled at the seafloor. The stars (x-axis labels) indicate significant differences determined by two-sample Wilcoxon tests.

For non-metazoans, the overall contribution of pelagic eDNA to the benthos increased gradually with depth across the water column, especially for the BBL samples (Fig. 4C). The proportions of pelagic eDNA sampled on nodules (9 % and 4 % for reads and ASVs, respectively) were over an order of magnitude higher than for metazoans. In contrast to metazoans, the percentage of ASVs per benthic sample that were simultaneously found in the pelagic was significantly higher in seamounts than in abyssal plain samples (t-tests; *p* < 0.001) (Fig. 5, Table S4). Significantly higher proportions of pelagic reads also could be observed within the 0-2 cm versus 3-5 cm sediment layer for the abyssal plain samples (Table S4, Fig. 5).

## 4 | Discussion

Biodiversity assessments in the deep ocean are essential prior to large-scale anthropogenic impacts, such as polymetallic nodule mining proposed for the Clarion Clipperton Zone (Smith et al., 2008a; Wedding et al., 2013, 2015). eDNA metabarcoding methods are increasingly being applied as a biomonitoring tool for ecosystem health in marine systems (e.g. Danovaro et al., 2016; Goodwin et al., 2017; Stat et al., 2017), but have seen limited application in the deep sea (e.g. Dell’Anno, Carugati, Corinaldesi, Riccioni, & Danovaro, 2015; Guardiola et al., 2015, 2016; Sinniger et al., 2016). Here we examine the effect of allochthonous pelagic DNA on eDNA surveys in abyssal habitats, and address several methodological issues that affect the optimal design of deep ocean eDNA biotic surveys.

Our results show that detrital pelagic eDNA represents a very small fraction of the diversity sampled in sediments and nodules at the abyssal seafloor (< 2 % of ASVs and their associated reads), and is unlikely to impede recovery of resident diversity in eDNA surveys of the deep ocean. Pelagic eDNA sampled in the abyss appears to derive from two distinct processes: (1) True legacy eDNA that is sourced from the high biomass upper ocean (epipelagic) and sinks as detrital particulate organic matter (POM) to the abyssal seafloor, and (2) eDNA sourced from resident populations in the bathy- and abyssopelagic ocean from organisms whose distributions are truncated at the seafloor. This interpretation derives from the fact that most of the epipelagic eDNA found in the benthos was observed in sediment samples, while meso- and bathypelagic eDNA found in the benthos was overwhelmingly present within the BBL seawater samples (Fig 4B-C). POM originating in the upper ocean is disaggregated and remineralized during its descent through the water column, declining exponentially with depth (Buesseler et al., 2007; Grabowski,Letelier, Laws, & Karl, 2019; Lutz, Caldeira, Dunbar, & Behrenfeld, 2007; Martin, Knauer, Karl, & Broenkow, 1987), with less than 2 % of primary production from the upper ocean typically reaching the abyssal seafloor (Smith et al., 2008b). This proportion is similar to that of the relative proportion of total epipelagic ASVs found in CCZ sediments (0.78 % and 1.29 % for metazoans and non-metazoans) and nodules (0.36 % and 2.65 % for metazoans and non-metazoans), suggesting that this material corresponds to true legacy eDNA and is acting as a tracer of POC flux to the abyssal seafloor. In BBL seawater samples, the proportion of epipelagic ASVs found at the seafloor was slightly higher (8.2 % and 5.8 % for metazoans and non-metazoans), and BBL pelagic reads were strongly augmented from material originating in the mesopelagic and below (up to 86 % and 72 % for metazoan and non-metazoans, respectively; Fig. 4C). For non-metazoans, both pelagic ASV and read proportions tended to increase with depth (Fig. 4C). Most of these deep-sourced sequences derived from hydrozoans, including siphonophore and trachymedusa families, such as Algamatidae, Rhopalonematidea and Prayidae, as well as radiolarians (Rhizaria). Hydrozoans can be either holoplanktonic or meroplanktonic, alternating between polypoid and medusoid forms. High relative abundance of hydrozoans, especially siphonophores, in deep pelagic habitats is well documented (e.g., Cartes, Fanelli, Lopez-Perez, & Lebrato, 2013; Robison, Sherlock, & Reisenbichler, 2010; Youngbluth, Sørnes, Hosia, & Stemmann, 2008), though there are also deep living benthic species (e.g., Rhodaliids live tethered to benthic substrates; O’Hara et al., 2016; Pugh, 1983), and their presence near the seafloor of the CCZ has been reported by Dahlgren et al. (2016). Rhizaria also occur in high relative abundance in the meso- and bathypelagic (Jing, Zhang, Li, Zhu, & Liu, 2018), and have been documented to be particle-associated in sinking POM material collected at abyssal depths in the Pacific (*see* Boeuf et al., 2019). It is likely that a high proportion of pelagic eDNA reads concurrently found in the BBL seawater samples originate from species whose distributions extend across a broad depth range, including the bathy- and abyssopelagic, and are truncated at the seafloor. We also found that non-metazoans and metazoans differed in the source of legacy eDNA to the seafloor, with detrital pelagic eDNA originating primarily from the epipelagic for metazoans, and across the water column for non-metazoans. Additionally, the 18S rRNA gene is highly conserved and has relatively low taxonomic resolution (Creer et al., 2016; Papadopoulou, Taberlet, & Zinger, 2015; Tanabe et al., 2016), and it is possible that close pelagic and benthic relatives are not differentiated by this marker. Based on the above, we infer that most metazoan pelagic eDNA found within sediment and nodules originates from detrital material sourced primarily in the epipelagic (‘true’ legacy eDNA), while pelagic eDNA found within the BBL mostly derives from residents whose distributions extend upward into the bathy- and abyssopelagic ocean.

Sedimentation rates over the CCZ are relatively low, estimated to range from 0.15 to 1.15 cm kyr^-1^ (Mewes et al., 2014; Müller & Mangini, 1980; Volz et al., 2018), and organic matter within the 0-5 cm layer may derive from sediment deposition over several thousand years (~ 4,300 to 33,300 yrs). Due to the low sedimentation rates and oxic conditions, most organic matter reaching the abyssal seafloor in the CCZ is remineralized within the upper few decimeters or meter of sediment (Burdige, 2007; Müller & Mangini, 1980; Volz et al., 2018). While eDNA in marine sediments can be preserved for several thousand years under certain conditions (Corinaldesi, Barucca, Luna, & Dell’Anno, 2011; Lejzerowicz et al., 2013), most sediment eDNA degrades over time scales of hours to months due to DNase activity and hydrolysis (depurination), or is taken up by microbial organisms as a nutrient source (Corinaldesi, Beolchini, Dell’Anno, & Bianche, 2008; Ibáñez de Aldecoa, Zafra, & González-Pastor, 2017; Nielsen,Johnsen, Bensasson, & Daffonchio, 2007). Dell’Anno & Danovaro (2005) measured extracellular DNA concentrations in deep-sea sediments and found exponential declines in eDNA in the top five centimeters, and estimate a mean eDNA residence time of 9.5 years for the top centimeter). In this study, using pelagic eDNA as a tracer to understand eDNA flux into and retention within the abyssal seafloor, we did not find evidence of increased pelagic eDNA degradation across a vertical profile within the sediment core for metazoans (Tables S4), since the great majority of samples had no pelagic ASVs. We did, however, find significant differences in sediment depth horizons for non-metazoans, with a higher proportion of pelagic reads in the 0-2 cm horizon in abyssal plains, and a trend toward a higher proportion of both pelagic ASVs and reads in the 0-2 cm layer across all abyssal habitats, as well as higher overall diversity within the 0-2 cm horizon (Figs. 5, S4). Note that a very small fraction of sediment eDNA was found to be pelagic in origin, and the overwhelming majority of eDNA sampled in the 0-5 cm layer derives from resident benthic organisms (historical and/or present assemblages).

Application of eDNA biotic surveys to monitor environmental impacts on the deep seafloor requires understanding the sampling effort required to adequately capture the diversity of deep benthic assemblages. Our results for abyssal sediments, polymetallic nodules and BBL seawater demonstrate highest ASV richness of both metazoans and non-metazoans at equivalent sampling coverage in sediments, with a substantial drop to the richness estimated for polymetallic nodules and BBL seawater (at base coverage, Fig. 2). Given prior studies demonstrating the importance of sample size to recovery of eukaryotic diversity (e.g., Nascimento et al. 2018), we targeted large sample volumes in comparison to many early eDNA studies, extracting DNA from 20 gr of sediment, 10 L of BBL seawater, and 1 gr of ground polymetallic nodule for each sample. For comparison, marine protist diversity on the continental shelf is typically sampled for eDNA using ~ 2 g of sediment (or less) in each sample (Chariton et al., 2015; Laroche et al., 2018; Pawlowski, Esling, Lejzerowicz, Cedhagen, & Wilding, 2014; Pochon et al., 2015), and metazoan surveys are often based on 1-2L of filtered seawater (e.g. Grey et al., 2018; Jeunen et al., 2019; Sigsgaard et al., 2017) or ~ 10 g per sample of the 3-5 top centimeter layer (deep sea examples, Guardiola et al., 2015, 2016; Sinniger et al., 2016). Despite relatively high sampling effort, including 71 sediment samples from 36 cores, 193 seawater samples from 12 CTD casts, and 50 samples from 10 polymetallic nodules, we are still under-sampling the diversity of these abyssal assemblages, with estimated sampling coverage for metazoans ranging from 40% to 74% study-wide across sample types. Our sampling coverage tended to be lower for metazoans than non-metazoans at equivalent sample size (e.g., in sediments 40% vs 64%), as expected since metazoans exhibit a higher degree of patchiness and because their detection relies in part on trace DNA recovery (Nascimento, Lallias, Bik, & Creer, 2018; Sinniger et al., 2016). For metazoans, diversity recovered from polymetallic nodules was approximately 6 times lower than in sediments at equivalent sampling coverage, and consequently, despite assessing only 5 g of material per nodule (~ 6 % of mean nodule weight) for a total of 10 nodules, sampling coverage was more complete study-wide and could be extrapolated to ~ 75 % of total community diversity. Nevertheless, the relatively low sampling coverage per nodule (~ 50 %) suggests that assessing additional material per nodule and increasing the number of sampled nodules would be beneficial. For seawater samples, metazoan ASV richness was higher at 5 mab within the BBL than any other depth sampled (at base coverage; Fig. 2C), with relatively low sampling coverage (49%, 57 %, BBL samples) despite filtering 10 L per sample (2 x 5 L). We therefore recommend increasing the number of replicates when sampling metazoan eDNA within the BBL, for example to > 30L (> 3 x 10L replicates) for sampling coverage at over 50 % of the community (Table 3).

With sampling coverage ranging from 64 % to 84 % for non-metazoans (all 3 substrate types), we infer that our sample sizes were adequate to capture the diversity of eukaryotic non-metazoans, although increasing sampling effort (# of samples) across a broader range of abyssal microhabitats would improve understanding of community structure and distribution of these assemblages. Sampling the abyssal seafloor is technically challenging and expensive relative to most marine environments, and as is commonly reported in other studies of abyssal fauna (e.g., De Smet et al. 2017), significant sampling effort is required to achieve high sampling completeness in particular for the sparse but highly diverse sediment community. Additionally, the detection probability of taxa is influenced by the amount of template DNA used for PCR amplification and in studies with low amount of template material, such as ancient DNA studies, a minimum of 8 PCR replicates have been proposed to increase detection probability (Ficetola et al., 2015). While we reduced stochasticity and increased detection probability by using a minimum of 3 ng of DNA per PCR and by pooling 2 PCR replicates per sample, further research evaluating the effect of PCR replication on deep sea benthic surveys would be valuable.

Overall, our results provide some guidance on designing eDNA biotic surveys for the deep ocean, as well as insight into processes that affect the transport and distribution of eDNA across the water column and abyss. Future deep-sea eDNA surveys should aim to process large amounts of material per sample (> 10 gr sediment, > 10 L seawater, with replication), and consider habitat heterogeneity at a range of spatial scales in the abyss. Finally, additional research is needed to evaluate how the oceanographic setting, including current velocities and sediment-transport regime, influence eDNA delivery to and retention within the abyssal benthos, and also to better understand the timescales for eDNA persistence in the abyss.

## 5 | Acknowledgements

We thank M. Church, E. Wear, other DeepCCZ scientists, and the captain and crew of R/V Kilo Moana for assistance at sea. We thank J. Saito from the Advanced Studies in Genomics, Proteomics, and Bioinformatics (ASGPB) core facility for laboratory support, and M. Gaither for early discussions on eDNA surveys in the deep ocean. Support for this project was provided by Pew Charitable Trusts (award UH30788, to PIs EG, CS), Gordon and Betty Moore Foundation award 5596 (lead PIs CS, J. Drazen), NOAA Ocean Exploration grant NA17OAR0110209 (lead PIs J. Drazen, CS), and from the University of Hawaii for ROV technical support at sea (lead PI CS). The views expressed are those of the author(s) and do not necessarily reflect the views of the Pew Charitable Trusts or other funding agencies.

## 6 | Author contributions

EG, OK, OL, and CS designed the study. EG, OK, and CS conducted field work and sampling at sea. OL generated the data and performed analyses, and OL wrote the manuscript with intellectual contributions from all co-authors. EG and CS provided grant and equipment support.

## 7 | Data Accessibility

Unprocessed sequences are accessible from the NCBI Sequence Read Archive (SRA) under accession numbers SRR9199590 to SRR9199869. Metadata for the samples are available in supplementary material.

