## Supplementary material for "From sea surface to seafloor: a benthic allochthonous eDNA survey for the abyssal ocean"

### 1 Supplementary Data

#### 1.1 Bioinformatic pipeline with Qiime2 (version 2018.11)

\# Directories

Database_fasta='Databases/SILVA_132_QIIME_release/silva_132_99_18S.fna'

Database_txt='Databases/SILVA_132_QIIME_release/18S_taxonomy_7_levels_NCBI.txt'

Metadata_file='Metadata.txt'

Outputs='Results_18S'

\# Quality and chimera filtering using DADA2

printf "\nStarting DADA2 process\n"

qiime dada2 denoise-paired \

--i-demultiplexed-seqs $Outputs/paired_end_demux.qza \

--p-trim-left-f 0 \

--p-trunc-len-f 260 \ # change to 240 for better resutls?

--p-trim-left-r 0 \

--p-trunc-len-r 235 \

--p-chimera-method consensus \

--p-min-fold-parent-over-abundance 1 \

--p-n-threads 0 \

--o-representative-sequences $Outputs/rep_seqs_dada2.qza \

--o-table $Outputs/table_dada2.qza \

--o-denoising-stats $Outputs/stats-dada2.qza \

--verbose

printf "\nCreating visualisation files of DADA2 outputs\n"

qiime feature-table summarize \

--i-table $Outputs/table_dada2.qza \

--o-visualization $Outputs/table_dada2.qzv \

--m-sample-metadata-file $Metadata_file

qiime feature-table tabulate-seqs \

--i-data $Outputs/rep_seqs_dada2.qza \

--o-visualization $Outputs/rep_seqs_dada2.qzv

qiime metadata tabulate \

--m-input-file $Outputs/stats-dada2.qza \

--o-visualization $Outputs/stats-dada2.qzv

\# Training a Naive Bayes classifier - assigning taxonomy from different databases

\## Importing the fasta and taxonomy files

printf "\nImporting reference database\n"

qiime tools import \

--type 'FeatureData[Sequence]' \

--input-path $Database_fasta \

--output-path $Outputs/database_seq.qza

qiime tools import \

--type 'FeatureData[Taxonomy]' \

--input-format HeaderlessTSVTaxonomyFormat \

--input-path $Database_txt \

--output-path $Outputs/database_taxonomy.qza

\# Extracting the sequence portion we have targeted

qiime feature-classifier extract-reads \

--i-sequences $Outputs/database_seq.qza \

--p-f-primer AGGGCAAKYCTGGTGCCAGC \

--p-r-primer GRCGGTATCTRATCGYCTT \

--o-reads $Outputs/database_seq.qza

\# Trainning the classifier

qiime feature-classifier fit-classifier-naive-bayes \

--i-reference-reads $Outputs/database_seq.qza \

--i-reference-taxonomy $Outputs/database_taxonomy.qza \

--o-classifier $Outputs/taxo_classifier.qza \

--verbose

\# Assigning taxonomy with the trainned classifier and looking at the results

qiime feature-classifier classify-sklearn \

--i-classifier $Outputs/taxo_classifier.qza \

--i-reads $Outputs/rep_seqs_dada2.qza \

--o-classification $Outputs/taxonomy.qza

\# Exporting tables

qiime tools export \

--input-path $Outputs/table_dada2.qza \

--output-path $Outputs/Data_analysis

biom convert -i $Outputs/Data_analysis/feature-table.biom \

-o $Outputs/Data_analysis/feature-table.txt --to-tsv

qiime tools export \

--input-path $Outputs/taxonomy.qza \

--output-path $Outputs/Data_analysis

qiime tools export \

--input-path $Outputs/rep_seqs_dada2.qza \

--output-path $Outputs/rep_set2.fna

#### 1.2 Seawater eDNA extraction protocol

In a pilot experiment of eDNA extraction methodologies, a modified protocol based on Pearl et al. (2008) and Shulse et al. (2017) provided the highest DNA yield and best purity measurement and was therefore used to process all the seawater samples using the following modified steps. First, filters were cut in half using sterile scissors and tweezers (soaked in 10% bleach solution for a minimum of 5 minutes and rinsed with ddH_2_O between samples). To mechanically disrupt cells, each half was placed in a 2 mL centrifuge tube containing 0.25 g and 1 g of sterile 0.1 mm and 1 mm silica beads respectively, and submitted to 3 consecutive thawing (incubated at 45°C for 90 sec) and freezing (flash-frozen in liquid nitrogen for 90 sec) cycles. 400 uL of solution AP1 was added to each tube, followed by 2 mins of bead-beating in a BioSpec Mini-beadbeater-16 (Oklahoma, USA) and 10 min of incubation at 65°C before being processed following the manufacturer’s protocol. Lysate from the halved filters was passed through the same QIAshredder column, and purified DNA eluted in 50 µL elution buffer. Due to low DNA concentration in the 5 and 50 mab samples (mean concentration of 0.573 ng/μL), replicates (2 x 5 L each) were pooled together and concentrated to ~ 1ng/μL with the DNA Clean & Concentrator kit (Zymo Research, California, USA) to obtain sufficient DNA for amplification.

### 2 Supplementary Figures and Tables

#### 2.1 Supplementary Figures

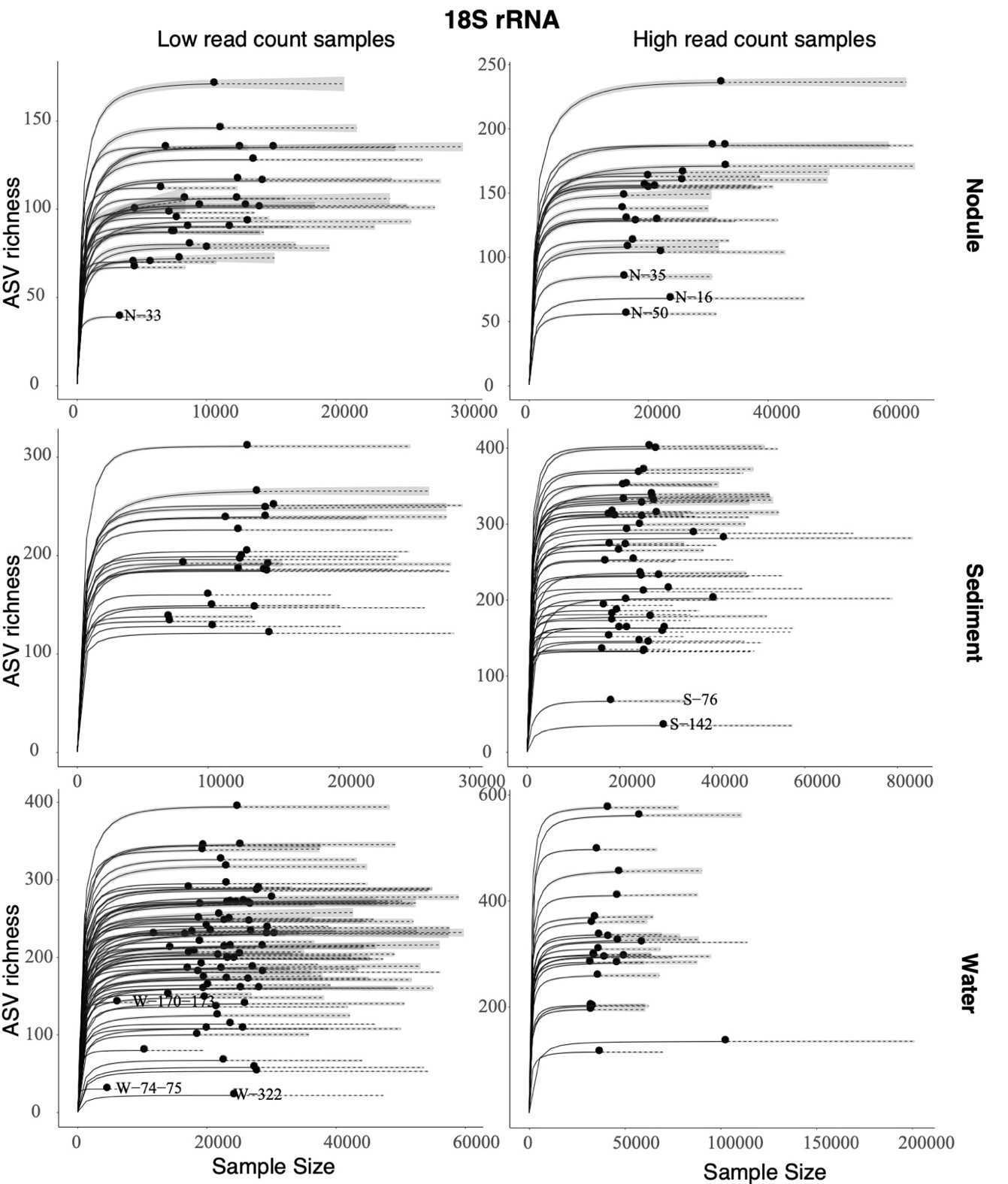

**Fig. S1.** Rarefaction curves of ASV richness across sequencing depth (Sample size) for all samples, plotted by sample type (polymetallic nodule, sediment, water). Sample names as labeled, with sampling metadata as reported in Table S1. Extrapolation values (dotted lines) are based on the Chao1 estimator. Grey shaded areas correspond to the 95 % confidence interval. Samples included in the low read count section of the figure (left side) had read counts below 15,000 reads for nodule and sediment, and below 25,000 for water.

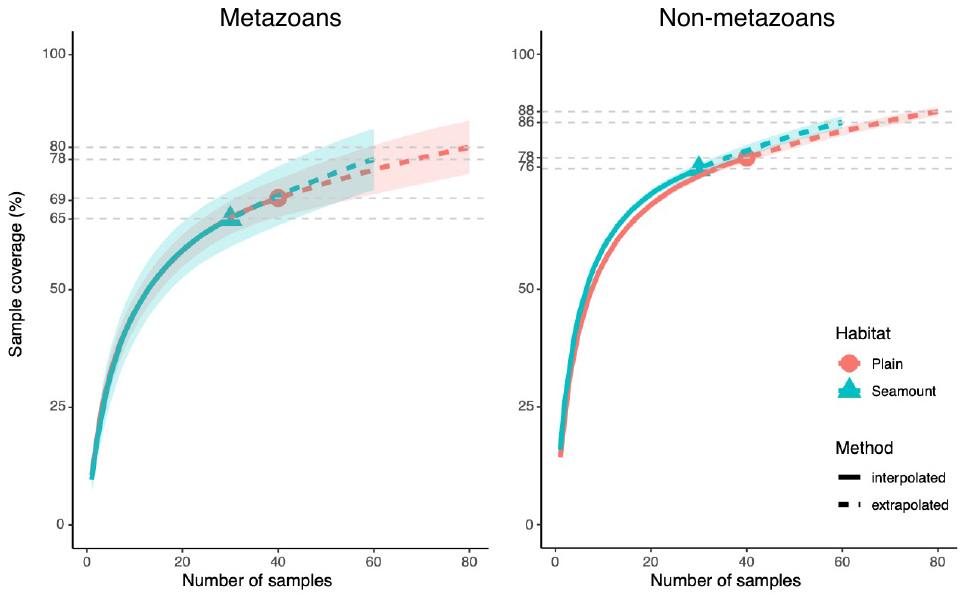

**Fig. S2.** Amplicon sequence variant (ASV) sampling coverage and richness for each habitat for both metazoans and non-metazoans. ASV richness was estimated using Chao2 (Chao & Jost 2012). Shaded, coloured areas indicate the 95 % confidence intervals obtained from a bootstrap method based on 200 replicates. Horizontal dotted grey lines (left, center panels) indicate maximum interpolation values for each habitat. Vertical dotted grey lines (right panels) indicate the value at base coverage, defined as the highest coverage value between minimum extrapolated values and maximum interpolated values.

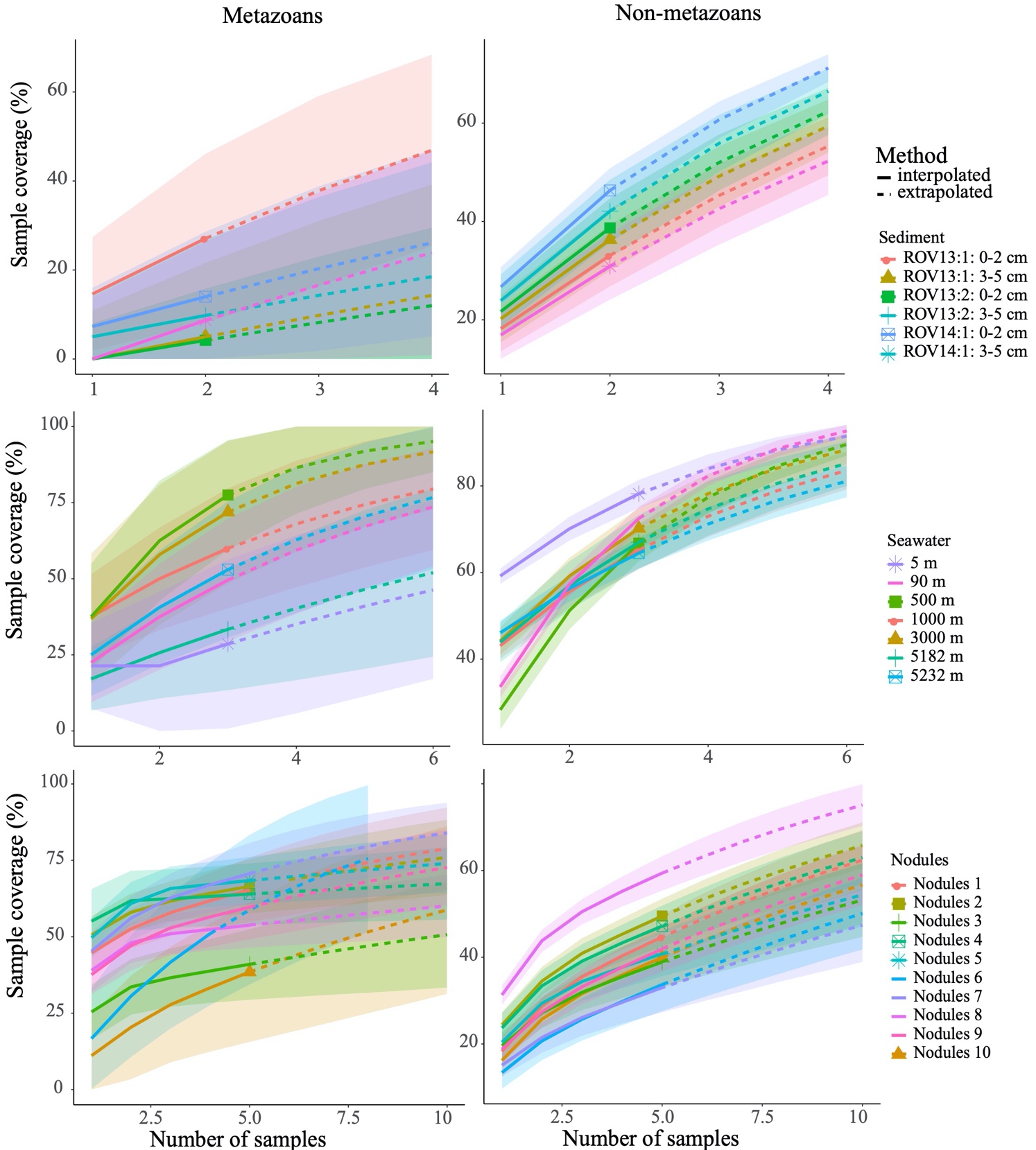

**Fig. S3.** Metazoan and non-metazoan (protist) sampling coverage per sample for each sample type. Shaded coloured areas indicate the 95 % confidence intervals obtained by a bootstrap method based on 200 replicates. Each of the five replicates per polymetallic nodule is composed of two nodule extractions of 0.5 g each.

**Fig. S4.** Venn diagrams displaying the number and proportion of unique and shared amplicon sequence variants (ASVs) between sediment layers and BBL depths for both metazoans and non-metazoans. The shading is proportional to the number of unique or shared ASVs, with darker shadings representing higher values.

#### 2.2 Supplementary Tables

**Table S4**: Two-sample Wilcoxon tests (two-sided) comparing the proportion of ASVs within benthic samples simultaneously found within the relevant APEI/Habitat combined pelagic samples between habitats and sediment layers. Significant p.values are in bold.

| **Dataset** | **Source\Habitat** | **Variable(s)** | **ASV/Reads** | **p.value** |
| --- | --- | --- | --- | --- |
| Metazoans | BBL | Habitat  (plains, seamounts) | ASVs | 0.340 |
|  |  |  | Reads | 0.982 |
|  | Sediment |  | ASVs | 0.905 |
|  |  |  | Reads | 0.973 |
|  | Plain | Sediment layer  (0-2 cm, 3-5 cm) | ASVs | 0.588 |
|  |  |  | Reads | 1 |
|  | Seamount |  | ASVs | 0.405 |
|  |  |  | Reads | 1 |
| Non-metazoans | BBL | Habitat  (plains, seamounts) | ASVs | **<0.001** |
|  |  |  | Reads | 0.770 |
|  | Sediment |  | ASVs | 0.517 |
|  |  |  | Reads | 0.095 |
|  | Plain | Sediment layer  (0-2 cm, 3-5 cm) | ASVs | 0.902 |
|  |  |  | Reads | **0.048** |
|  | Seamount |  | ASVs | 0.936 |
|  |  |  | Reads | 0.180 |
